# Flagellar synchronization is a simple alternative to cell cycle synchronization for ciliary and flagellar studies

**DOI:** 10.1101/089185

**Authors:** Soumita Dutta, Prachee Avasthi

## Abstract

The unicellular green alga, *Chlamydomonas reinhardtii*, is an ideal model organism for studies of ciliary function and assembly. In assays for biological and biochemical effects of various factors on flagellar structure and function, synchronous culture is advantageous for minimizing variability. Here, we have characterized a method in which 100% synchronization is achieved with respect to flagellar length but not with respect to the cell cycle. The method requires inducing flagellar regeneration by amputation of the entire cell population and limiting regeneration time, which results in a maximally homogeneous distribution of flagellar length at three hours post-amputation. We find that time-limiting new protein synthesis during flagellar synchronization limits variability in the unassembled pool of limiting flagellar protein and variability in flagellar length without scaling the cell volume range. We also find that long and short flagella mutants that regenerate normally require longer and shorter synchronization times, respectively. By minimizing flagellar length variability using a simple method requiring only hours and no changes in media, flagellar synchronization facilitates the detection of small changes in flagellar length resulting from both chemical and genetic perturbations in *Chlamydomonas*. This method increases our ability to probe the basic biology of ciliary size regulation and related disease etiologies.

## Introduction

The unicellular, biflagellate alga *Chlamydomonas reinhardtii* is extensively used as a model organism for studying fundamental processes such as photosynthesis, cell motility, cell signaling, cell-cell recognition and regulation of ciliary assembly-disassembly (1). This organism offers many advantages for molecular and biochemical studies of eukaryotic flagella as their flagellar structure and function is well conserved (2). *Chlamydomonas* cells can be chemically or mechanically induced to shed their flagella (termed deflagellation). After amputation, they can regenerate flagella rapidly to pre-deflagellation lengths within 2 hours. Flagellar assembly and disassembly are precisely controlled throughout cell-cycle progression and cell division (3, 4). During cell division flagella are disassembled naturally. Flagellar resorption starts at the pre-prophase stage and continues about 30 minutes prior to mitotic cell division (5). New flagella begin to form in the daughter cell after division (6, 7). During the sexual cycle, flagella begin to resorb a few hours after the fusion of gametes and proceed gradually as in vegetative growth (8).

As cell division plays a critical role in flagellar growth and resorption, cultures with a heterogeneous population of cells in different divisional stages vary in their flagellar length. In contrast, synchronous cultures, which contain cells that are in the same growth stage, have a comparatively homogeneous distribution of flagellar length. Thus, synchronous cultures provide advantages over non-synchronous cells for studying cellular morphology and the effects of various chemical or genetic perturbations on flagellar length.

A wide range of physical and chemical methods have been applied to achieve synchronization for different cells or tissue types. Synchronization of bacteria can be carried out by single or multiple changes of temperature or light, single or multiple cycles of nutritional starvations, cell-cycle inhibitor block and size selection by filtration or centrifugation (9–12). Fission yeast can be synchronized either by separating a subpopulation from an asynchronous culture using specialized centrifugation or by selecting cells from a lactose gradient (13). Temperature sensitive cell cycle mutations or inhibitors are also used to block the cell cycle at different stages of growth, which allows cells to grow synchronously upon withdrawal of the block (14). Common methods for mammalian cell cycle synchronization are either inhibition of DNA replication (15) or inhibition of mitotic spindle formation using different chemical inhibitors (16–18). Non-chemical methods for cell cycle synchronization include amino acid and serum starvation (19). Cells can also be mechanically separated by physical methods such as flow cytometry, mitotic shake-off or counter-current centrifugal elutriation (18). Hypoxic and hyperthermic shock have been used to synchronize the ciliate *Tetrahymena pyriformis* (20). Photosynthetic algal cells are typically exposed to alternative light/dark cycles for synchronization (21, 22).

In *Chlamydomonas reinhardtii*, like other photoautotrophic cells, the most common method used for cell synchronization is alternating light/dark cycles (12 hour/12 hour or 14 hour/10 hour) in minimal medium (23, 24), though other methods such as periodic hypothermic conditions (25), selection by size (26) or variable wavelengths of light (27) have been applied. Synchronization can also be achieved by incubating *Chlamydomonas reinhardtii* cells in low nitrogen medium for at least 15 hours (28). *Chlamydomonas* cells undergo gametogenesis upon nitrogen deprivation. After induced gametogenesis, culture contains mostly new-born cells with smaller sizes (29–31). During light-dark synchronization (L-D synchronization), cells can grow during the light phase to many times their original size (32). In the dark phase, cells can undergo consecutive divisions to produce 2, 4, 8, 16 or even 32 daughter cells depending on the cell size (33). In *Chlamydomonas reinhardtii*, cells divide in the middle of the dark cycle (23) whereas in *Chlamydomonas moewussi* cells, the division occurs during the late phase of darkness (34). Although cell division is restricted to each dark phase, the starting time of individual cell divisions varies from cell to cell. Thus, consecutive cell divisions take place throughout several hours of the dark period. As a result, the cells are always partially asynchronous in their division at any point in time (8). In addition, cultures are maximally synchronized only after 3^rd^ iteration of light-dark cycling since some population of the cells divide during 1^st^ and 2^nd^ iterations of the light phase (23). Different factors such as light duration and intensity, temperature and culture density also have an effect on the degree of homogeneity (32, 35). For example, in *Chlamydomonas eugametos*, L-D synchronization can be achieved only if the culture is static without aeration (36).

While synchronization can be optimized, it is not possible to synchronize entire cell populations by any of the methods or techniques described above (37). Traditional L-D synchronization or nitrogen starvation methods can only make partially synchronized *Chlamydomonas* cultures. As cells are not truly synchronized using these methods, high variabilities of flagellar length are still observed within the population. If the culture contains too much heterogeneity, it can be difficult to detect effects of flagellar length perturbations. As synchronized cells are ideal for assaying length related flagellar kinetics, here we have outlined and characterized a method in which 100% of cells are synchronized with respect to their flagellar length but not synchronized with respect to the cell cycle. We tested the utility of this method when evaluating chemical and genetic perturbations to flagellar length. Finally, we probed the basis of flagellar synchronization by probing the synthesized but unassembled pool of flagellar protein.

## Results

### Flagellar length synchronization narrows the steady state flagellar length distribution

Synchronous culture provides a better way to address cell cycle and related flagellar kinetics. However, cell cycle synchronization methods provide only partial synchronization and thus show high variance in flagellar length. To obtain 100% flagellar length (F-L) synchronization, we exploit an inherent property of *Chlamydomonas*, which is their ability to regenerate the flagella after amputation (38). We first performed different synchronization methods and then compared their steady-state flagellar lengths to non-synchronized cells (Fig.1a, Table S1). For F-L synchronization, we tested different regeneration time durations following deflagellation to determine the time point at which flagellar length variability is minimized. We found that the pre-deflagellation flagellar length distribution was broad, which was expected as our starting culture was non-synchronous and contained a heterogeneous population of cells (Fig. 1b, red). After deflagellation, all flagella started to grow synchronously but the length distribution still remained broad at 2 and 2.5 hours, when some cells were still in the 8-9 μm range and did not reach their original length (Fig. 1b, light green). The distribution narrowed and became maximally homogeneous at 3 hours (Fig. 1b, green, Table S2). However, the length-distribution remained narrow only for a short time, expanding again within 30 minutes and increasing with time (Fig. 1b, dark green). This timing was highly reproducible and Fig. S1 is the combined result of three independent experiments. Based on the standard deviations of these distributions (Fig. 1b and Fig. S1, lower panel), we selected 3 hours regeneration time post-deflagellation as the flagellar length synchronization time and the time at which to initiate further experiments. Likewise, after determining conditions of minimal variability for each synchronization method, we measured the steady state flagellar lengths for comparative analysis. The results revealed that the mean flagellar lengths at steady state were almost equivalent for all methods (Table S1). As expected, non-synchronous cells had larger variability compared with all other synchronized cells (Fig. 1a, red). The F-L synchronization method shows a remarkably narrow spread of measurements around the mean and the smallest variability of length across all synchronization methods (Fig. 1a, green and standard deviation bar graph below, Table S1). In contrast, conventional L-D synchronization and M-N synchronization have comparatively wider distributions (Fig. 1a, blue and purple respectively, Table S1). Our findings suggest that the F-L synchronization is the most effective method for achieving maximum flagellar-length homogeneity.

**Figure 1:**
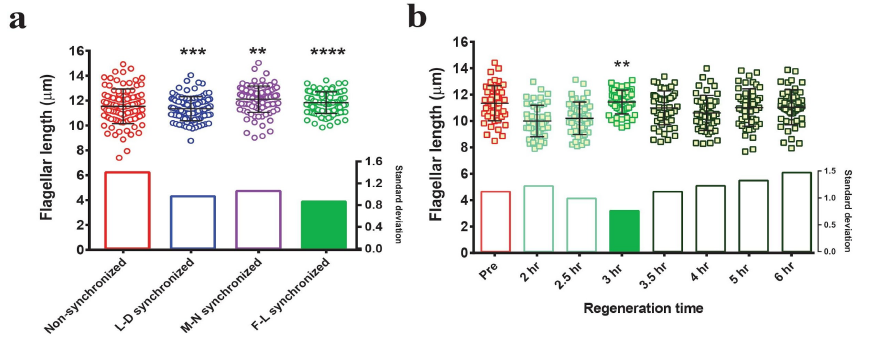
F-L synchronization narrows the flagellar length distribution compared to other synchronization methods. **a)** Distribution of steady state flagellar length after different synchronization methods. Non-synchronous cells were used as a control and steady state flagellar lengths were measured after each synchronization method as described in the text. (N=100/each). F-test was performed for comparing variance and two-tailed p value was considered. `Asterisks indicate significant differences (**** p≤0.0001, *** p≤0.001, ** p≤0.01)`. Standard deviations of each distribution are shown below the individually plotted values. **b)** Wild type flagellar-length distribution at various time intervals during the regeneration after amputation. Pre-deflagellation non-synchronous cells (pre) are shown in red. Regeneration was carried out for indicated times after deflagellation by pH shock (green). Lighter and darker green indicates before and after the time of F-L synchronization respectively. (N=50/each). F-test was performed for comparing variance (control= pre-deflagellation non-synchronous cells). Bonferroni corrected α_altered_= 0.007. Asterisk indicates significant difference below α_altered_ (**p=0.004). Standard deviations are expressed as bar graphs in the bottom half of panels. The filled SD bar represents the condition with the smallest variance.

### Increased effects of chemical perturbations using F-L synchronization

Flagellar length can be perturbed chemically. If the perturbation has a small effect on flagellar length, high variance in the system may mask observed phenotypes. Our observations demonstrate that F-L synchronized cells have reduced variance in flagellar length compared to other methods (Fig. 1a). Therefore, we tested the effects of several known flagellar length altering agents after reducing variability in the initial population through F-L synchronization and compared the results with other synchronization methods. When flagellar shortening was induced with 3-isobutyl-1-methylxanthine (IBMX) (39), latrunculin B (LatB) (40), and sodium pyrophosphate (NaPPi) (41), we observed more severe shortening in F-L synchronized cells compared to all the other methods (Fig. 2a-c, green, Table S3). Only L-D synchronization, which is a more time consuming synchronization method, demonstrates length reduction comparable with F-L synchronized cells (Fig. 2a-c, blue, Table S3). Effects on flagellar length are the most extreme in the case of NaPPi mediated length resorption (Fig. 2c). After the treatment, flagellar length distribution was significantly narrowed in F-L synchronized cells compared to the others and produced a more dramatic shortening of the mean flagellar length (Fig. 2c, Table S3). L-D synchronized cells showed a reduced effect (~41% shortening) compared to F-L synchronized cells (~46% shortening) (Fig. 2c, lower panel). In addition to flagella shortening compounds, we also tested the effects of lithium chloride (LiCl), which is known to lengthen flagella (6). The flagella were longest in F-L synchronized cells after treatment of LiCl. While the variance and % change in mean flagellar length were comparable between L-D and F-L synchronized cells, each of the LiCl treated F-L synchronized cells had flagellar length longer than 13.5 μm with an average of 17 μm (Fig. 2d, green, Table S3). Broad distributions of lengths were observed in both nonsynchronized and M-N synchronized cells ranging from 9 μm to 20 μm (Fig. 2d, red and purple respectively). As a result, the effect on flagellar length was less apparent. Taken together, all of these data demonstrate that the effect of each chemical is more prominent and detectable in F-L synchronized cells when the variance in starting flagellar distribution is minimized.

**Figure 2:**
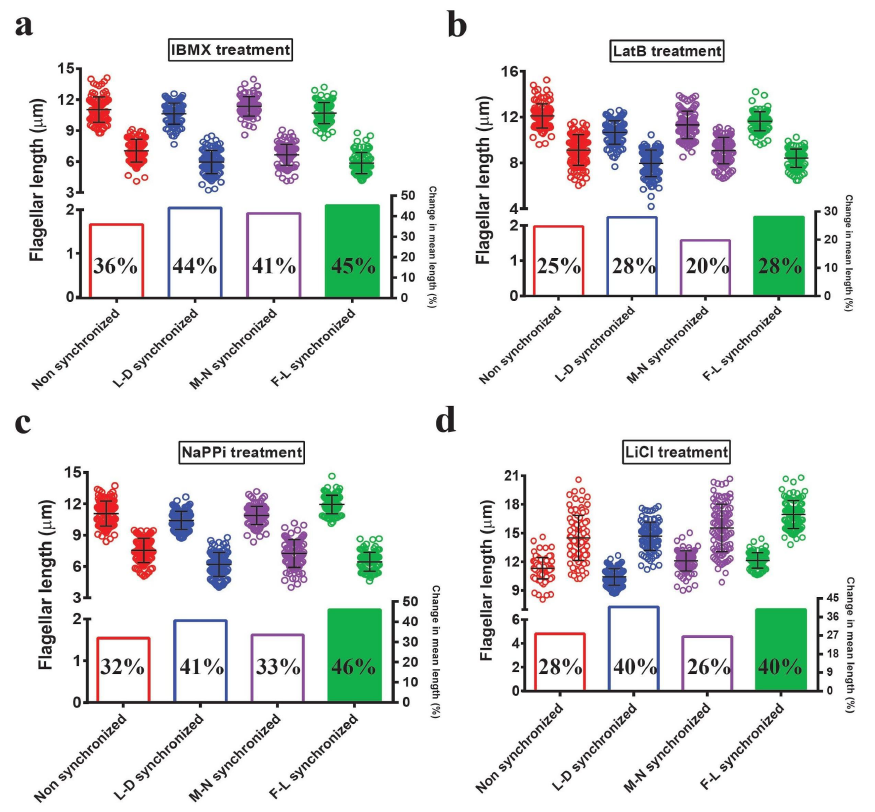
Effects of length-altering chemical treatments are more apparent after F-L synchronization. Flagella were measured from each cell after incubation with appropriate concentration of different chemicals for 90 minutes: **a)** 0.4 mM IBMX, **b)** 10 μM LatB, **c)** 10 mM NaPPi and **d)** 25 mM LiCl. For each pair: non-synchronized (red), L-D synchronized (blue), M-N synchronized (purple) and F-L synchronized (green), the first and second distributions represent control and treated respectively. N=100 cells (one flagellum per cell). Bars are mean and standard deviation (top half of panel). In each case, the percentage change in mean flagellar length is shown below individual plotted values.

### Synchronization time varies in flagellar length mutants

In *Chlamydomonas*, flagellar length mutants with both long and short flagella have been isolated (42–44). Length distributions are reportedly wider in flagellar length mutants than wild type cells (43). Therefore, we asked if F-L synchronization would increase our ability to detect length differences in these populations. Some long flagella mutants have defective regeneration kinetics after amputation (43) and therefore are not suitable for F-L synchronization. However, other mutants have regeneration kinetics comparable to wild type cells (43). We first considered the *lf4-7* mutant cells which have the longest flagella of all identified long flagella (lf) mutants but can regenerate their flagella with wild type regeneration kinetics after amputation (44). As expected, prior to deflagellation, the initial population of *lf4-7* cells were distributed very widely (12 μm to 28 μm) (Fig. 3a, upper and lower panel, Table S4). Following amputation, flagellar length variabilities were reduced at the 2 and 3 hour time points, but mean lengths had not yet achieved pre-deflagellation lengths (Fig. 3a). We found that at least four hours were required to regenerate flagella to their pre-deflagellation length. Like wild type cells, flagella also took extra time after reaching their original length (6 hours of regeneration in this case) to become homogeneously distributed (Fig. 3a). Also like wild type cells, a narrow distribution could be maintained for only a short period of time. For a mutant with short-flagella, *shf1-253* (42), flagella reached their pre-deflagellation lengths within 1.5 hours following deflagellation, but took an additional one hour (2.5 hours) to distribute more narrowly (Fig. 3b, upper and lower panel, Table S4). Finally, we studied a mutant with only slightly longer flagella than wild type (45), *cnk2-1*. Like other length-mutants, these cells regenerated to normal length after 3.5 hours following amputation but achieved the narrowest flagellar length distribution at 5 hours of regeneration (Fig. 3c, upper and lower panel, Table S4). These findings suggest that all cells exhibit their tightest flagellar length distribution sometime after they initially reach pre-deflagellation mean lengths. Therefore, mutants with longer flagella take longer and mutants with short flagella take less time to achieve their most narrow flagellar length distribution. Ideally, the optimal time for flagellar synchronization should be adjusted for individual strains.

**Figure 3:**
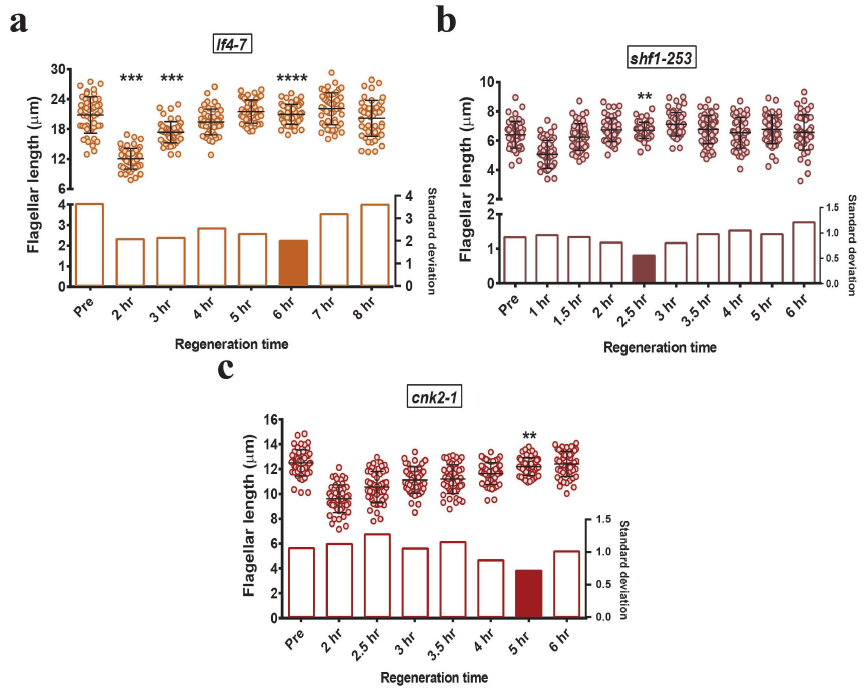
F-L synchronization time varies in different length mutants. For each mutant, distributions of flagellar length during regeneration are shown. **a)** long-flagella mutant, *lf4-7*, **b)** short-flagella mutant *shf1-253* and **c)** mildly long flagella mutant *cnk2-1*. ‘Pre’ represents the steady state length of the mutant pre-deflagellation. Bars are mean and standard deviation (top half of each panel). Standard deviations are represented by bar graphs in the lower half of each figure and the filled bar corresponds to the synchronization time for each mutant on the basis of minimal standard deviation. N=50/ea. F-test was performed for comparing variance with pre-deflagellation non-synchronous controls. For (a) and (c), Bonferroni corrected α_altered_= 0.007 and asterisks indicate significant differences below α_altered_ (****p= 0.00005, ***p= 0.0002, **p= 0.004). For (b), Bonferroni corrected α_altered_= 0.0055 and asterisks indicate significant differences below α_altered_ (**p=0.0008).

### F-L synchronization may mask important outliers

Some flagellar length mutants have a mean flagellar length comparable to wild type cells but have a positively skewed distribution that includes small numbers of extremely long flagella (43). As, F-L synchronization reduces length variance, we asked if the informative long outliers would be lost after minimizing variability and thereby decrease our ability to appropriately phenotype this class of mutant. In order to test this, we chose two long-flagella mutants *lf2-5* and *lf3-2*, which were able to regenerate their flagella normally and had a large number of flagella in the wild type range (43). When we induced regeneration for these two mutants up to 8 hours, we found that F-L synchronization time for both *lf2-5* and *lf3-2* was 4 hours. The flagellar length distributions of *lf3-2* cells demonstrated that synchronized cells had a narrow distribution with flagella no longer than 20 μm and no shorter than 10 μm, which was expected. The synchronized distribution has a negative kurtosis (−0.3738) i.e, a distribution with short tails compared to the non-synchronized distribution which has a positive kurtosis (+0.0358) with relatively long tails. The average length of synchronized compared to the non-synchronized population was not changed significantly (Fig. 4a). The mode changed from 12.5 to 16 with the number of cells containing 13 μm to 16 μm long flagella increasing remarkably in case of F-L synchronized cells (Fig. 4a). Therefore, for *lf3-2*, F-L synchronization did not affect our ability to identify a mutant phenotype despite eliminating long outliers. In the case of *lf2-5*, F-L synchronization also removed outliers from the mutant population (Fig. 4b) but this time the average flagellar length between non-synchronized and synchronized cells differed markedly (Fig. 4c) by left-shifting the distribution. The mode changed from 16 to 13 and *lf2-5* showed average flagellar length value ~14 μm after F-L synchronization, which sometimes is seen in wild type populations (Fig. S2). Also, non-synchronized and synchronized kurtosis values (−0.6492 and −0.6753 respectively) are not significantly different. While F-L synchronization of *lf3-2* and *lf2-5* retains our ability to discriminate between wild type and mutant phenotypes by reducing the variance (Fig. 4c), the two mutants behave differently with respect to changes in descriptive statistics. Therefore, losing outliers during F-L synchronization has the potential obscure important information when evaluating genetic perturbations. We therefore recommend testing both synchronized and non-synchronized cells when characterizing new mutants.

**Figure 4:**
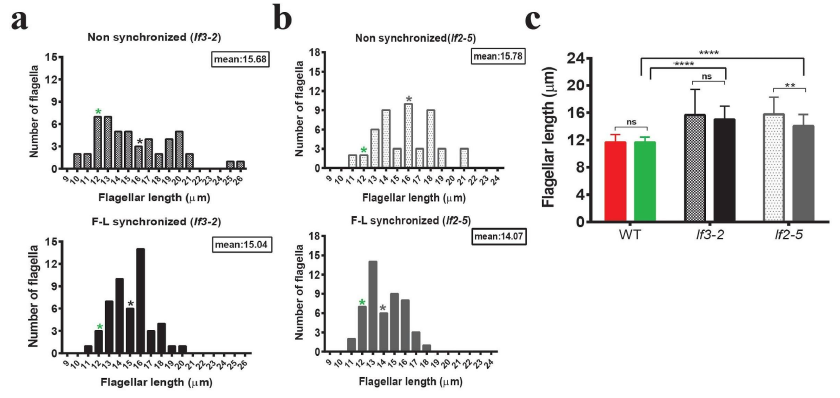
Distributions of flagellar length in long-flagella mutants before and after F-L synchronization. **a)** *lf3-2* and **b)** *lf2-5*. The upper panel of figure (a) and (b) represent the distribution of flagellar length in non-synchronous cells. Cells were then deflagellated by acidic shock and regenerated for 4 hours for F-L synchronization. Lower panel of figure (a) and (b) correspond to the distribution of flagellar length after F-L synchronization. N=50/ea (zeros excluded). The mean flagellar length of each mutant is marked by asterisk. Green asterisk indicates the mean flagellar length observed in wild type controls. The mean flagellar length of wild type and mutant cells were compared using Bonferroni’s post hoc test and the data were represented in figure **(c)**. The first and second bars within each pair in figure (c) represent mean flagellar length of non-synchronized and synchronized cells respectively. Mann-Whitney U test was performed for comparing two means in a pair and Bonferroni’s post hoc test was performed for multiple comparisons. Asterisks indicate significant differences (**** p≤0.0001, ** p≤0.01, ns p>0.05).

### Flagellar length variability is related to precursor pool variability

As flagellar synchronization time is highly reproducible within wild type populations, we hypothesized that there might be an internal regulator which is responsible for the narrow distribution pattern after 3 hours of regeneration. *Chlamydomonas* cells have a synthesized pool of unassembled flagellar proteins or at least a pre-existing pool of some protein that limits the rate of flagellar assembly (termed precursor pool). The size of this pool is sufficient to assemble flagella to half of their normal length if new protein synthesis is inhibited (46). Limiting-precursor models of flagellar length control have been previously considered but flagellar length appears to be maintained independent of pool size or concentration (42). However, completely blocking new protein synthesis can limit flagellar length so we asked if putting constraints on protein synthesis and incorporation might narrow the resulting flagellar length distribution. In this case, reduced variability during F-L synchronization would be due to synchronizing flagellar protein synthesis through deflagellation and time-limiting flagellar protein incorporation. In order to test our hypothesis, we determined the variance of synthesized precursor pool size after different synchronization methods by deflagellating cells and allowing them to regenerate in the presence of cycloheximide (46). This allows existing flagellar protein to be incorporated into flagella but prevents the synthesis of new protein. In these experiments, flagellar length is a proxy for the amount of limiting protein available for incorporation into flagella without protein synthesis. Precursor pool size is therefore reported in units of “μm flagellar length.” To evaluate the relationship between flagellar variability and precursor pool variability, we compared flagella that had undergone 3 hours of regeneration (tightest distribution), and two other time points with increased variability: 2 hours and 5 hours (Fig.1b and Fig. S1). We performed the same for nonsynchronized, L-D synchronized and M-N synchronized cells. As expected, all cells after synchronization but prior to deflagellation for cycloheximide treatment recapitulated our previous findings (Fig. S3, Table S5). When we compared variance of precursor pools after different regeneration time intervals, we found a narrow distribution of pool size at 3 hours of regeneration (green) but not in 2 hours (light green) or in 5 hours (dark green) (Fig. 5a, lower half of 5b, Table S5). Non-synchronous cells have a precursor pool distribution comparable to 2 and 5 hours of regeneration in F-L synchronized cells (Fig. 5a, red). For all F-L synchronized cells, we observed that the variance of the precursor pool (Fig. 5b, lower panel) echoes the post-synchronization length variance (Fig. 5b, upper panel). We also saw that both L-D and M-N synchronized cells showed reduced variance in cytoplasmic precursor pool of flagellar proteins (Fig. 5a, blue and purple respectively, Table S5). Because both L-D and M-N synchronization are cell cycle synchronization methods, we asked next if the narrow precursor pool variability in these cells correlated with the cell size distribution and if we were circumventing the cell size dependence of the flagellar precursor pool by time-limiting protein synthesis and incorporation.

**Figure 5:**
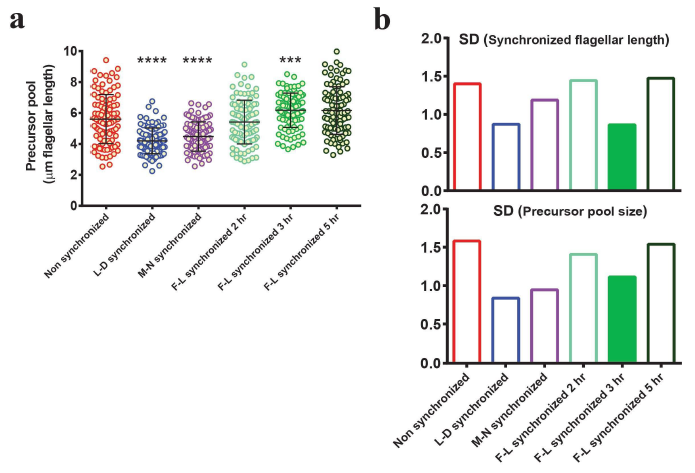
Relationship between flagellar length and precursor pool distribution. **a)** Flagellar precursor pool distributions (as determined by regeneration in cycloheximide after different synchronization methods). Non-synchronous (red), L-D synchronized (blue), M-N synchronized (purple), F-L synchronized for 2 hours (light green), F-L synchronized for 3 hours (green) and F-L synchronized for 5 hours (dark green) are compared. F-L synchronization for 3 hours shows minimal variability among F-L synchronized cells. N=100/ea. F test was performed for comparing variance (control= non-synchronous cells). Asterisks indicate significant differences (**** p≤0.0001, *** p≤0.001). Bars are mean and standard deviation. **b)** Post-synchronization standard deviations (top panel) and corresponding precursor pool standard deviations (bottom panel) of each distribution are represented by bars with matching colours. Cell cycle synchronized cells show the smallest variability in precursor pool distribution (blue, bottom panel). Of F-L synchronized cells (green), 3 hours synchronization shows the smallest variability in precursor pool distribution (filled bar, bottom panel).

### Flagellar length synchronization changes the relationship between cell size and precursor pool size

It is reported that there is no simple relationship between cell size, flagellar length and precursor pool size (47). However, we observed that relative variability of flagellar length across synchronization methods is preserved when considering the variability in pool size (Fig. 5b). Given the general scaling of protein quantity with cell size (48–52), we tested the relationship between cell size and precursor pool size to better understand the factors influencing precursor pool and flagellar length variance across synchronization methods. We regenerated the flagella for 2 hours in the presence of cycloheximide after different synchronization methods and measured flagellar length to determine the pre-existing precursor pool as before. This time, we also measured the corresponding cell volume. In nonsynchronized cells, cell volumes had a broad distribution (~100 μm^3^ to 900 μm^3^) as did the precursor pool size distribution (~2.5 μm to ~9 μm flagellar length), with a significant correlation between cell size and precursor pool size (r= 0.73, two tailed p<0.0001) (Fig. 6a). Expectedly, we found that both in L-D and M-N synchronized cells, cell volumes were very restricted, ranging from ~100 μm^3^ to ~400 μm^3^ (Fig. 6b and 6c respectively) as they are cell cycle synchronized. Since smaller cells generally produce less protein (48), restricted cell volumes of cell cycle synchronized populations also limited the protein precursor pool within a very narrow range (~4.5 μm flagellar length) (Fig. 6b and 6c). On the other hand, like non-synchronous cells, F-L synchronized cells had a large cell size range (~100 μm^3^ to 900 μm^3^). However, unlike non-synchronous cells with a precursor pool size ranging ~6.5μm flagellar length (Fig.6a), F-L synchronized cells had a narrow precursor pool range (~5 μm flagellar length) more comparable to the precursor pool size in cell cycle synchronized cells (Fig. 6d). While a correlation between cell size and precursor pool size was still maintained in F-L synchronized cells (r= 0.71, two tailed p<0.0001), the slopes of the regression lines for non-synchronous and F-L synchronized cells were statistically different (p=0.0012) (53–55) (Fig. 6e, red and green line respectively). In other words, the relationship between cell size and available precursor pool is altered upon F-L synchronization. Presumably, F-L synchronization can limit the precursor pool sizes by limiting the amount of time the precursor pool can accumulate without the need to limit pool size by restricting cell size through cell cycle synchronization.

**Figure 6:**
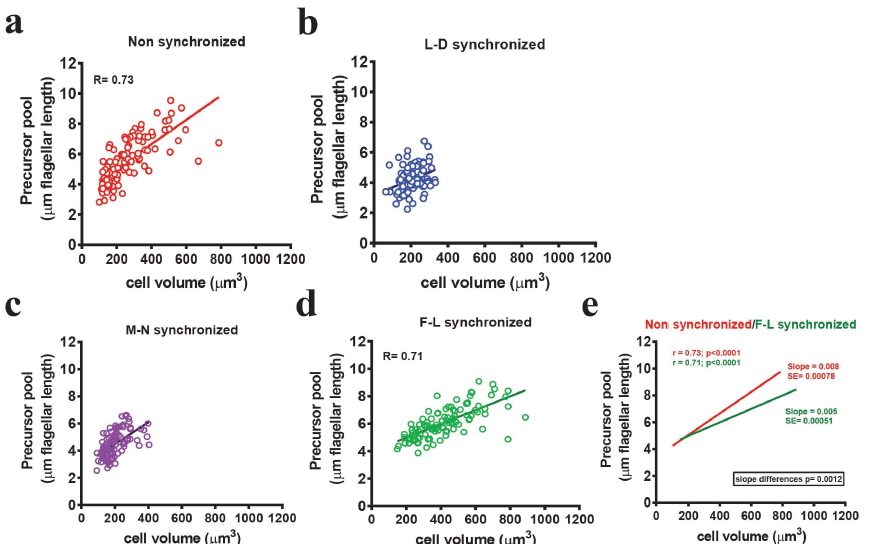
F-L synchronization changes the relationship between the precursor pool size and cell size. Flagellar length in cycloheximide, which corresponds to the precursor pool, is plotted along the Y axis. Matching cell volume is plotted along the X axis; Values are plotted for **(a)** non-synchronized (red), **(b)** L-D synchronized (blue), **(c)** M-N synchronized (purple) and **(d)** F-L synchronized (green) populations. N=100. Non-synchronized cells show a significant correlation between cell size and precursor pool size (r=0.73). L-D and M-N synchronized cells have narrow cell volume and precursor pool ranges. F-L synchronized cells show a narrow precursor pool range without limiting cell volume. **(e)** Relation between non-synchronous and F-L synchronized cells. The straight lines represent the best fitted lines through the data point which were drawn after linear regression (red: non-synchronous cells and green: F-L synchronized cells). SE= standard error of the slopes. The slope became smaller in case of F-L synchronized cells which is significantly different from non-synchronous cells (p=0.0012).

## Discussion

Here we have shown a powerful new approach to understanding ciliary length-related biology by characterizing a synchronization method that minimizes flagellar length variability. It is well established that flagellar length variability can be controlled by restricting the cell size (the basis of cell cycle synchronization). Our data suggests that the precursor pool size (existing pool of flagellar protein not assembled into flagella) is also related to the cell size. Limiting flagellar protein is not currently considered a major factor controlling flagellar length because flagellar proteins in *Chlamydomonas* clearly exceed the amount assembled into flagella, unassembled flagellar precursor pool size does not correlate with the flagellar length during assembly and the length of flagella do not appear to be strongly dependent upon the number of flagella in mutants with variable flagellar number (41, 47, 56). However, while the precursor pool size is correlated with cell size, cell cycle synchronization methods severely limit both cell size and precursor pool variability. We further found that we can circumvent cell size restriction for minimizing flagellar length variability by limiting the amount of time the flagellar precursor pool can accumulate and incorporate into flagella. It is well known that mammalian cell ciliary studies often initiate ciliogenesis by serum starvation of confluent cells (5, 19). By standardizing plating density and limiting serum starvation time prior to subsequent experimentation (thereby limiting the time window of assembly), the F-L synchronization method may be applicable to studies in mammalian cells.

Assembly of full length flagella requires pre-existing precursor pool, *de novo* synthesis of flagellar precursor proteins and also incorporation of those proteins into the flagellar structure (57). The expression of genes encoding ciliary proteins is dramatically up-regulated after flagellar amputation to replenish the precursor pool and to provide proteins required for flagellar assembly (58). We propose a model for F-L synchronization (Fig.7) where F-L synchronization via deflagellation works by stimulating a highly regulated program of gene expression and flagellar protein incorporation so that all cells can regenerate their flagella synchronously regardless of their divisional phase. Synchronization is then achieved by limiting the time window (Fig. 7, red dotted line) that the cells are allowed to regenerate their flagella (three hours). The combination of simultaneous induction of the regeneration program along with imposing a restriction of the amount of protein synthesis and incorporation result in a tighter length distribution pattern.

**Figure 7:**
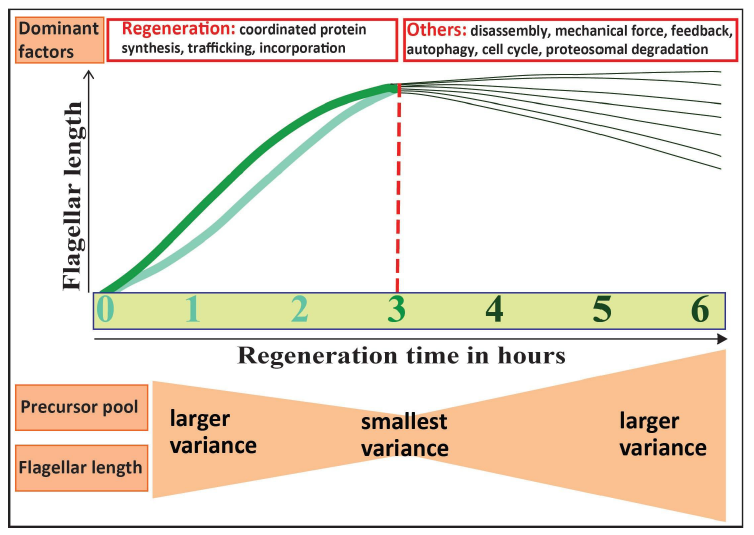
Proposed model of flagellar length synchronization. Thick light green and green lines represent slow growing and fast growing flagella respectively. Red dotted line denotes the three hour post-deflagellation time at which flagellar length variability is minimized. Initially regeneration is the dominant factor. With time, other listed factors may contribute, resulting in larger flagellar length variability.

If time limiting protein synthesis and incorporation results in a narrow distribution of flagellar lengths, why is there an increased variability of flagellar lengths at an earlier flagellar synchronization interval (2 hours)? Rates of flagellar regeneration vary from cell to cell; some flagella are fast-growing (Fig. 7, green line) and attain their original length within 2 hours of amputation while slow-growing flagella (Fig. 7, light green line) can reach only 80% of their length within that period. We saw in measurements of unassembled flagellar protein (Fig. 5b, lower panel, Table S5) that cells that have undergone 2 hours of F-L synchronization have a smaller precursor pool (mean: 5.4μm flagellar length) compared to 3 hours (mean: 6.2μm flagellar length). The slow growth of some cells at 2 hours post-deflagellation may therefore be due to reduced protein synthesis and accumulation. We propose that, as we extend the regeneration time beyond two hours, slow-growing flagella finally catch up their original length and fast-growing flagella approach their maximum length by reducing their assembly rate and reaching equilibrium with continuous disassembly (Fig. 7). To confirm this, we would need data on the individual cell level rather than at the population level, which will be performed in future studies by trapping individual motile cells in a micro-fluidic chamber (59).

While all cells must initiate a regeneration program upon deflagellation, with increasing time, the deflagellation-induced protein synthesis and incorporation program (which decreases as a function of time and flagellar length) may be overcome by other regulating factors such as disassembly, mechanical force, proteosomal degradation, feedback control or autophagy (60–64) (Fig. 7, dark green lines). Also, when the regeneration program no longer drives flagellar length after 3 hours, cell cycle regulation may dominate, resulting in the heterogeneous flagellar length and precursor pool distributions seen in nonsynchronized cells.

In addition to maximizing our ability to detect effects in inhibitor studies, we observed that F-L synchronization can be readily applied to genetically perturbed length mutants to reduce their length heterogeneity. All mutants will have different genetic defects and we showed several mutants that respond differently to synchronization, highlighting that both synchronized and non-synchronized cells should be tested for proper phenotyping of newly identified mutants. Interestingly, based on the synchronization time alone, we were able to discriminate long flagella mutants from wild type cells regardless of their mean length. In other words, when F-L synchronization eliminated important outliers and reduced the ability to discriminate on the basis of mean flagellar length, still showed a flagellar synchronization profile more similar to long flagella mutants (4-5 hours synchronization time) than wild type cells (3 hours synchronization time). This suggests flagellar synchronization time itself can be a useful phenotyping parameter.

Currently, the most commonly used method of reducing flagellar length variability in *Chlamydomonas* is cell cycle synchronization using L-D cycling. However, in L-D synchronized cells, natural variance of cells (37) prevents 100% synchronization of flagellar length. Using F-L synchronization, we can synchronize 100% of the population through deflagellation and produce a homogeneous distribution of length by 3 hours. Conventional L-D synchronization, by contrast, requires at least three days achieving comparable levels of homogeneity. F-L synchronization does not require a dark chamber with automated light switching. Moreover, the entire experiment can be performed in rich medium like TAP instead of minimal medium which is very sensitive to changes in pH. This facilitates the use of inhibitors that would otherwise dramatically affect pH of the medium. F-L synchronization demonstrated equivalent or stronger effects of length-altering chemicals on flagellar length compared to L-D synchronization demonstrating its utility in addition to its uniformity and simplicity.

The study presented here facilitates identification of ciliary length-related defects (65–68) by increasing our ability to detect small changes in cilia size but more broadly helps us understand factors affecting ciliary length regulation. By inducing a fully synchronous cellular program (regeneration) that temporarily dominates multiple other factors to minimize flagellar heterogeneity, F-L synchronization also has the strong potential to benefit studies of ciliary motility or ciliary signaling.

## Materials and methods

### Strains and length altering chemical treatments

*Chlamydomonas reinhardtii* wild type137c mt+ [CC125], *lf4-7* mt− [CC4534], *shf1-253* mt− [CC2348], *cnk2-1* [CC4689], *lf2-5* mt− [CC2287] and *lf3-2* mt− [CC2289] were obtained from the *Chlamydomonas* Resource Center at the University of Minnesota. All chemicals were purchased from Sigma (St. Louis, MO) and final concentrations of 0.4 mM IBMX, 10 mM NaPPi, 10 μM LatB, 25 mM LiCl and 10 μg ml^−1^ cycloheximide were used. Compounds were diluted to the indicated doses either with Tris-Acetate-Phosphate (TAP) medium or with 100% dimethyl sulphoxide (DMSO). For the chemical treatment, 1 or 2 ml of cells were treated with indicated concentration of chemicals with indicated controls and placed on a rotator for 90 minutes or 120 minutes as indicated in the text.

### Culture conditions and synchronization methods

All cells were maintained on TAP plates containing 1.5% agar (Difco laboratories, Detroit, MI) (24). For liquid cultures, cells were inoculated from TAP plates less than two weeks old.

#### Non-synchronous culture

For non-synchronous culture, cells were grown in liquid TAP medium for 24 hours on a culture rotator drum at 25 °C under continuous illumination with an LED LumiBar with independent red and blue light control (LumiGrow, Inc.).

#### Light/Dark synchronization (L-D synchronization)

Cells were inoculated in minimal medium (M1 medium) from the TAP plates and kept in light for 12 hours and then dark for 12 hours alternating at 25 °C for at least three days. After each light/dark cycle (12 hour/12 hour), cultures were diluted to 2 × 10^5^ cells ml^−1^ with fresh M1 medium. On the 4^th^ day, after growing at light phase for 5 hours, cultures were immediately transferred to TAP medium prior to chemical treatment.

#### Synchronization by nitrogen starvation (M-N synchronization)

M-N synchronization was attained by inducing gametogenesis in nitrogen free minimal medium (i.e, M minus N) for 18-20 hours in continuous light at 25 °C under a LumiBar. These cells were then transferred to TAP medium for 4 hours prior to chemical treatment.

#### Flagellar length synchronization (F-L synchronization)

For F-L synchronization, *Chlamydomonas* cells were grown in liquid TAP medium and then induced to regenerate flagella after acid-mediated flagellar excision (69). 60 μl of 0.5 N acetic acid was added to 1 ml of cells for deflagellation (pH= 4.5). Immediately after 45 seconds, 70 μl of 0.5 N KOH was added to neutralize the medium which ultimately induced the flagellar regeneration. Wild type cells were grown for 3 hours for flagella-regeneration under continuous illumination with a LumiBar on a rotator drum.

### Flagellar length and cell volume measurement

For flagella measurements, cells were fixed in 1% glutaraldehyde and kept at 4 °C. Cells were then centrifuged at 1000 × g for 1 minute and mounted between a glass slide and coverslip. Imaging was performed using Zeiss Axioscope differential interference contrast (DIC) microscope with a 40X objective lens and a Zeiss Axiocam 105 color camera. Flagellar length measurements were done by line segments and spline fitting using Image J software (NIH, USA). All flagella in a particular field were considered and at least 50 flagella were measured at each time point. For cell size determination, cell volumes were calculated using the ellipsoid equation 4/3π [L/2][W/2]^2^, where L is cell length and W is cell width (70). Flagellar length distributions and cell volumes were plotted using GraphPad Prism software version 6 (GraphPad, USA).

### Flagellar precursor pool determination

For flagellar precursor pool determination, cells were allowed to regenerate their flagella in the presence of 10 μg ml^−1^ cycloheximide following deflagellation (46). Cells from different synchronization methods were induced to regenerate flagella after acidic shock and then returned to neutral pH by adding KOH as described above and subjected to cycloheximide treatment immediately. For precursor pool determination in F-L synchronized cells, cells were allowed to regenerate their flagella for 2 hours, 3 hours and 5 hours after the 1^st^ deflagellation and then subjected to a 2^nd^ deflagellation prior to cycloheximide treatment. For all cases, cells were centrifuged at 1000 × g for 2 minutes after neutralizing and then resuspended in TAP medium containing cycloheximide. Cells were placed on a rotator for 120 minutes and flagellar length measurements were carried out to determine the amount of unassembled limiting flagellar protein.

### Statistical analysis

Statistical analysis was performed using GraphPad Prism software version 6 and excel-2010. Descriptive statistics were expressed as mean and standard deviation (SD). F test was performed in excel 2010 for comparing the variance of the dataset and to determine the p values (two tailed). Non parametric Mann-Whitney U test was performed for comparing two means. One-way analysis of variance (ANOVA) and Bonferroni’s post hoc test were performed for multiple comparisons and to determine p values. For all datasets, p <0.05 was considered statistically significant. However, Bonferroni’s correction was applied when multiple pair-wise tests were performed on a single set of data and p values were adjusted accordingly. Frequency distribution and column statistics were carried out for determining mode and kurtosis. For r value, SE and slope determination, we performed correlation and linear regression (least square fit). We also determined the difference between the two slopes using available online software (53, 54).

## Acknowledgements

We thank members of the Avasthi Lab and members of the Cilium Interest Group for their help and valuable comments. We thank Sumedha Gunewardena and Joshua Habiger for their helpful discussions in statistical analysis. This work was supported by NIH P20 GM104936 and NIH P20 GM103418.

## Author Contributions

Experiments were designed by SD and PA. All experiments and flagellar/cell measurements were carried out by SD. Manuscript was written by SD and PA.

